# Ectopic bone formation and systemic bone loss in a transmembrane TNF-driven model of human spondyloarthritis

**DOI:** 10.1101/2020.05.14.095588

**Authors:** Eleni Christodoulou-Vafeiadou, Christina Geka, Lydia Ntari, Ksanthi Kranidioti, Eleni Argyropoulou, Florian Meier, Marietta Armaka, Iordanis Mourouzis, Constantinos Pantos, Maritina Rouchota, George Loudos, Maria C Denis, Niki Karagianni, George Kollias

## Abstract

**Background:** The transmembrane-TNF transgenic mouse, TgA86, has been shown to develop spontaneously peripheral arthritis with signs of axial involvement. To assess similarity to human spondyloarthritis we performed detailed characterization of the axial, peripheral and comorbid pathologies of this model.

**Methods:** TgA86 bone pathologies were assessed at different ages using CT imaging of the spine, tail vertebrae and hind limbs and characterized in detail by histopathological and immunohistochemical analysis. Cardiac function was examined by echocardiography and electrocardiography and bone structural parameters by µCT analysis. The response of TgA86 mice to either early or late anti-TNF treatment was evaluated clinically, histopathologically and by µCT analysis.

**Results:** TgA86 mice developed with 100% penetrance spontaneous axial and peripheral pathology which progressed with time and manifested as reduced body weight and body length, kyphosis, tail bendings as well as swollen and distorted hind joints. Whole body CT analysis at advanced ages revealed bone erosions of sacral and caudal vertebrae as well as of sacroiliac joints and hind limps, and also, new ectopic bone formation and eventually vertebral fusion. The pathology of these mice highly resembled that of SpA patients, as it evolved through an early inflammatory phase, evident as enthesitis and synovitis in the affected joints, characterized by mesenchymal cell accumulation and neutrophilic infiltration. Subsequently, regression of inflammation was accompanied by ectopic bone formation, leading to ankylosis. In addition, both systemic bone loss and comorbid heart valve pathology were evident. Importantly, early anti-TNF treatment, similar to clinical treatment protocols, significantly reduced the inflammatory phase of both the axial and peripheral pathology of TgA86 mice.

**Conclusions:** The TgA86 mice develop a spontaneous peripheral and axial biphasic pathology accompanied by comorbid heart valvular dysfunction and osteoporosis, overall faithfully reproducing the progression of pathognomonic features of human spondyloarthritis. Therefore, the TgA86 mouse represents a valuable model for deciphering the pathogenic mechanisms of spondyloarthritis and for assessing the efficacy of human therapeutics targeting different phases of the disease.

## Background

Spondyloarthritis (SpA) describes a group of chronic inflammatory disorders that have been initially classified as clinically distinct and included ankylosing spondylitis (AS), psoriatic arthritis (PsA), reactive arthritis (ReA), inflammatory bowel disease (IBD)-associated arthritis and more (1,2). More recently, these pathologies have been unified based on common and overlapping features including inflammation of peripheral joints, the spine and/or sacroiliac joints as well as structural damage dominated by bone destruction and progressive bone formation eventually evolving into axial and peripheral joint ankylosis (1,3–5). Additional features include extraarticular manifestations such as psoriasis, uveitis, IBD, aortitis and osteoporosis (6,7).

The development of SpA has been linked to several factors, with a focus on genetic factors such as HLA-B27 (2,6) but also on the contribution of several pro-inflammatory cytokines (8,9) that are central to the transformation and perpetuation of disease including TNF-α, IL23, IL17, IL1β etc.(10). Despite the associations with pathogenic factors, the understanding of SpA pathogenesis remains incomplete and therapeutic approaches are targeting symptoms rather than the cause of the pathology.

Therefore, animal models of SpA are of high value as they can significantly contribute to the better understanding of the disease (11), addressing important questions such as the identification of the pathogenic factors that underlie disease causality and contribute to disease mechanisms responsible for the development of specific features of the pathology. A faithful mouse model of SpA should ideally reproduce both the progression, as well as the full complexity of the human pathology including main disease features such as inflammation of the spine and sacroiliac joints progressing to bone formation and ankylosis, peripheral joint arthritis, enthesitis and dactylitis, accompanied by one or more relevant extraarticular manifestations of the disease.

The pro-inflammatory cytokine Tumor Necrosis Factor alpha (TNF-α) is a key pathogenic factor involved in the development of SpA, with evidence stemming primarily from the significant therapeutic effect of TNF-blocking agents that, when administered early on in the development of the disease, can alleviate SpA symptoms (12–15). It has been previously shown that overexpression of mouse TNF through the deletion of AU-rich elements in its 3’UTR, leads to the development of destructive polyarthritis, sacroiliitis, enthesitis, Crohn’s-like IBD and aortic valve inflammation, however without evidence of new bone formation (16,17). Similarly, mouse models overexpressing human TNF (18,19) have been shown to develop spontaneous arthritis also missing however evidence of new bone formation, which is the main feature of SpA.

Interestingly, the TgA86 transgenic mouse model overexpressing specifically mouse transmembrane TNF (tmTNF) signaling through both TNFR1 and TNFR2 receptors has been previously shown to develop both peripheral and axial clinical manifestations (20), thus prompting us to further explore whether this model is faithfully reproducing key pathologic features of human SpA. Through computed tomography (CT) imaging and detailed histopathological analysis we study here in great detail the features of the axial and peripheral pathology of the TgA86 mice and align them to those of human SpA. We show that the TgA86 model closely reproduces major aspects of human SpA including the development of spondylitis, sacroiliitis, enthesitis, and bone destruction gradually leading to activation of new bone formation mechanisms and ankylosis, while accompanied by comorbid heart valve pathology and osteoporosis. We propose that the TgA86 is an animal model faithfully reproducing the complexity of human SpA, a statement further supported by the finding that, in line with clinical findings in human patients, early anti-TNF treatment leads to the alleviation of its SpA manifestations. Overall, we propose that this transmembrane TNF dependent mouse model is suitable for the study of SpA pathogenic mechanisms and for the *in vivo* efficacy evaluation of human therapeutics targeting different aspects of SpA pathology either at the early or later stages of the disease.

## Methods

### Mice

The TgA86 mice overexpressing tmTNF have been previously described (20). Briefly, these mice carry a 3.2 kb genomic DNA fragment containing 5’-regulatory sequences and a muTNFΔ1-12 mutated gene linked to the 3’-UTR of the human β-globin gene. Mice used in this work were heterozygous for the transgene and were maintained on a CBA×C57BL/6J genetic background in Biomedcode’s animal facility, under SPF conditions. All mice experimentation was performed according to National legislation. The wellbeing of all mice was regularly monitored and mice were euthanized when needed according to animal welfare guidelines. For the initial phenotyping experiments, the termination time point was set to a maximum of 40 weeks of age.

For the evaluation of the effect of anti-TNF treatment, groups of 6-10 age- and sex-matched TgA86 mice were treated subcutaneously either with saline or Etanercept 30mg/kg (Amgen/Pfizer) thrice weekly from 2.5 (early) or 9 weeks (late) up to 20 weeks of age.

Mice were monitored on a weekly basis to assess the clinical signs of peripheral and axial pathology. A group of age- and sex-matched wild type (wt) littermates was used as a control group. At the end of the treatment period animals were sacrificed, and tails, hearts, ankle and sacroiliac joints were collected and processed for further analysis.

### Clinical Scoring

The *in vivo* peripheral and axial pathology of the TgA86 mice was monitored on a weekly basis from 2.5 to 20 weeks of age. A peripheral pathology scoring system was developed to assess the severity of joint swelling, digit/limb deformation and grip strength using a scale from 0-2 (Additional file 1; Table S1). Similarly, an axial pathology scoring system was developed, that assessed severity on a scale from 0-3 based on evaluation of tail bendings and stiffness (Additional file 1; Table S2).

### Body length measurements, CT and µCT imaging

Live photos of animals at 6, 10 and 15 weeks of age were acquired under standard conditions and they were used to assess body length, from nose to tail tip, using the ImageJ processing software. CT imaging was performed using the x-CUBE (Molecubes), with no further need of a contrast agent. Mice were lightly anaesthetized with isoflurane and acquisition was performed using a high-resolution and Multi Rotation protocol at 50kVp. For whole body acquisitions, an Iterative Image Space Reconstruction Algorithm (ISRA) with 200µm voxel size was used. For the local scans of specific regions, the reconstruction was done with 100µm voxel size. Images were exported and post-processed on VivoQuant software, version 4.0 (Invicro, Boston) and further analysis was also performed with RadiAnt DICOM viewer (64-bit).

μCT imaging was performed using the Sky Scan 1172 μCT imaging system (Bruker). Tail and femur specimens were fixed in 4% aqueous formaldehyde solution overnight and scanned at 50kV, 100μA, 5W, with aluminum filter 0.5mm, zoom 12, medium camera (2000×1332), rotation step 0.4-0.8. 3D reconstruction images of the caudal vertebra C6 were generated using NRecon 1.6.8.4 software (Bruker). C6 vertebra length was measured on the shadow projection of tail scannings using the CTAn V1.14.4.1. software (Bruker). Structural indices of femur trabecular bone (BV/TV, TbN, TbSp, Conn Den) were calculated using the CTAn analysis program.

### Histological analysis and Immunohistochemistry

Ankle joints, tails, sacroiliac joints and hearts were harvested and fixed in 4% formalin. Ankle joints, tails and sacroiliac joints were decalcified in EDTA-decalcification buffer, embedded in paraffin and cut into 4μm sections. Paraffin sections were stained with Hematoxylin/Eosin (H&E), Safranin O (s/o), tartrate-resistant acid phosphatase (TRAP) (SIGMA) or with specific antibodies in combination with the Vectastain Elite ABC HRP and the Vectastain DAB kits (Vector Laboratories). The antibodies used for immunochistochemistry included Gr1 (MCA2387GA; BioRad), Periostin (ab14041; Abcam), Osteopontin1 (OPN) (AF808; R&D Systems), B220 (553084; BD Pharmigen) and Vimentin (ab92547; Abcam). Images were acquired with Leica DM2500 microscope equipped with Leica SFL4000 camera (Leica Microsystems). Histopathological evaluation of peripheral and axial inflammation was performed on H&E stained sections, according to the scoring criteria outlined in Additional file 1; Tables S3 and S4.

The assessment of vertebral inflammation was performed on H&E tail sections focusing on 4-6 consecutive caudal vertebrae of each mouse (6-10 mice at each age of 10, 20 and 40 weeks of age) and was based on the scoring system described in Additional file 1; Table S3. Inflammation scores were classified in three different groups: low (score 0-1), medium (score >1 and ≤2) and high (score >2 and ≤3) inflammation; percentages represent the ratio of the number of inflamed vertebrae of a given severity to the total number of vertebrae scored for each age group.

Osteoclasts were counted in TRAPs stained sections of TgA86 tails, focusing on the site of inflammation (enthesis) and adjacent area of 4 successive vertebrae of each mouse (2-3 mice of 10, 20 and 40 weeks of age).

New bone formation evaluation was performed on s/o stained sections, according to a scoring system ranging from 0-3 described in Additional file 1; Table S5.

### Aortic Valvular thickness measurement

Three independent measurements of the widest portion of the aortic valve leaflets were taken from 2 consecutive H&E-stained transverse heart sections using the ImageJ software (NIH).

### Echocardiography and Electrocardiogram

Echocardiography and electrocardiogram assessment was performed in 40-weeks of age animals, as previously described (21).

### Statistical analysis

Data are presented as mean±SEM, and Student’s t test was used for the evaluation of statistical significance, with P values <0.05 being considered statistically significant.

Analysis was performed using the GraphPad Prism V.6.

## Results

### TgA86 tmTNF transgenic mice model key features of human SpA

TgA86 mice have been previously described to develop peripheral arthritis with signs of axial involvement (11,20). Here we analyze in greater detail these mice with a primary focus on the characterization of their axial pathology, thus assessing its similarities to human SpA.

TgA86 mice presented with a significantly reduced body weight and body length compared to wild type (wt) mice of the same age (Fig. 1a, b). As early as 3 weeks of age, they developed clinical signs of arthritis in all four limbs, which manifested as joint swelling, digit/limb deformation and joint stiffness ultimately resulting in loss of grip strength (Fig. 1c). Axial pathology was clinically evident as tail bending and eventually tail stiffness also accompanied by hyperkyphosis better evident by CT imaging (Fig. 1c, d). All pathological features worsened over time.

**Figure 1.**
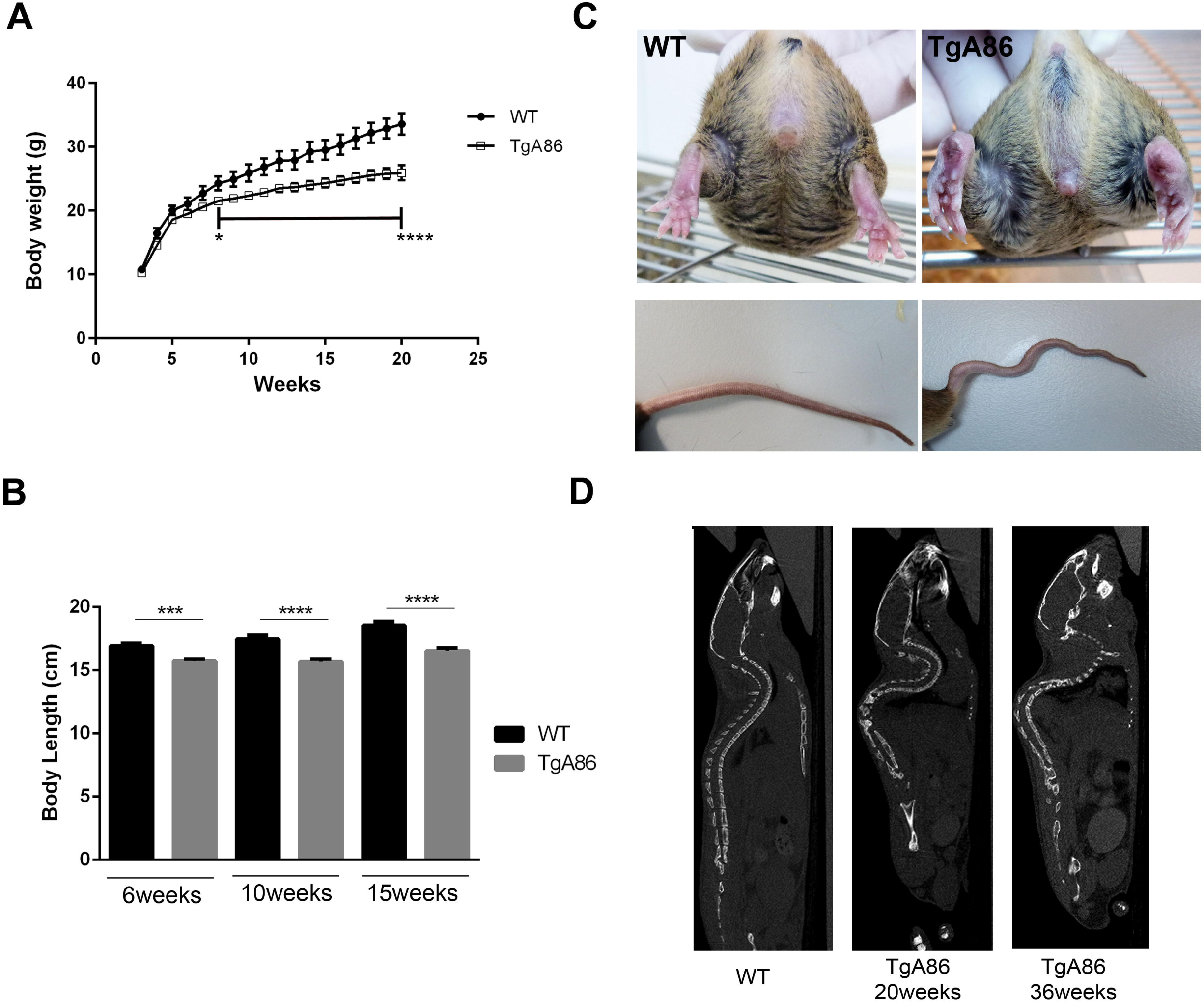
Phenotypic characteristics of the TgA86 mouse model. **(A, B)** TgA86 transgenic mice, starting from an early age, exhibit significantly reduced body weight **(A)** and body length **(B)** compared to wt littermates (data are presented as mean±SEM; ***P<0.0002; ****P<0.0001). **(C)** Representative photos at 20 weeks of age, highlight joint swelling and digit distortion in the hind limbs of TgA86 mice (upper panel) as well as characteristic bendings of their tails (lower panel). **(D)** Sagittal views of whole-body CT images show the hyperkyphosis observed in 20-week-old and, more severely, in 36-week-old TgA86 mice.

To better characterize the TgA86 axial and peripheral pathology, we performed CT imaging of 40-week-old TgA86 mice and wt littermates focusing on sites known to be important also in human SpA pathology, such as the spine, including the cervical, thoracic, lumbar, sacral and caudal segments, as well as the sacroiliac joints. In addition, we also examined the hind limbs to gain a better grasp of the peripheral pathology. Interestingly, CT imaging revealed previously undetected pathological features both in the spine, especially at the sacral vertebrae, and the sacroiliac joints. More specifically, the frontal view of the TgA86 sacrum revealed structural changes in sacral vertebrae S3/S4 involving loss of their intervertebral space and fusion of the vertebrae, evident in 83% of the transgenic mice examined (Fig. 2a; red arrow). The side view of sacrum, also showed loss of the intervertebral space (Fig. 2b; red arrow) and fusion of the spinous processes, evident in 80% of the TgA86 mice examined (Fig. 2b; yellow arrow). Moreover, the superior parts of the sacral ala presented with bilateral structural damages in TgA86 mice (Fig. 2a; yellow boxes).

**Figure 2.**
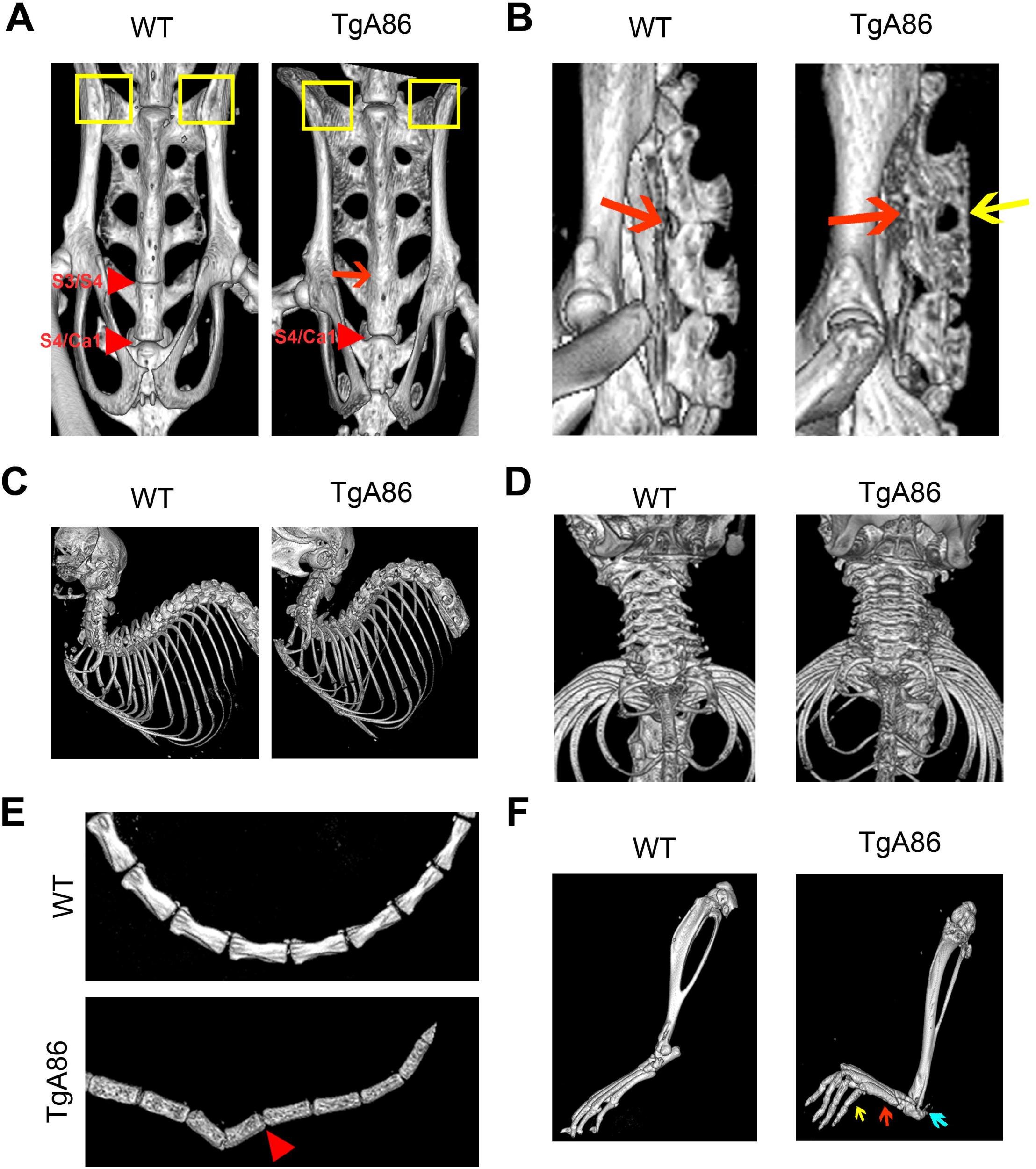
CT imaging reveals major changes in the spine, tail, sacroiliac and hind joints of TgA86 mice. 40-week-old TgA86 mice were imaged by CT. **(A)** Frontal view of the sacrum of TgA86 mice demonstrates fusion of the sacral vertebrae 3 and 4 (red arrow) as well as structural changes at the sacral ala (yellow box). **(B)** Fusion of the bodies (red arrow) as well as of the spinous processes (yellow arrow) of the sacral vertebrae is also seen in the sagittal view of the sacrum. **(C, D)** Sagittal **(C)** and frontal **(D)** views of the cervical and thoracic areas of the spine of TgA86 mice reveal hyperkyphosis and compressed vertebrae respectively. **(E)** CT imaging of the TgA86 tails reveals vertebral bridging and fusion (red arrowhead) while **(F)** CT imaging of the TgA86 hind limbs shows thickened metatarsal bones (red arrow) and phalanges (yellow arrow), as well as structural changes at the posterior calcaneus (blue arrow) and the knee joints.

CT imaging of the upper body revealed extensive hyperkyphosis (Fig. 2c). Cervical vertebrae of TgA86 mice were found to be compressed with reduced intervertebral spaces (Fig. 2d), while lumbar as well as thoracic vertebrae did not show major abnormalities.

CT imaging of the caudal area, where axial pathology of TgA86 mice was originally observed (20), revealed significant changes of the vertebrae structure including loss of their bar bell shape and roughening of their surface accompanied in many cases by vertebrae bridging and fusion (Fig. 2e; red arrow).

CT imaging of the peripheral joints revealed thickening of the metatarsal bones (Fig. 2f; red arrow) and phalanges (Fig. 2f; yellow arrow), roughening of their surface as well as structural deformation of the posterior calcaneus at the site of Achilles entheseal insertion (Fig. 2f; blue arrow), while knee and ankle joints presented also with roughened surfaces and deformed tibia and femur heads (Fig. 2f).

To assess alignment of the TgA86 phenotype, with human SpA pathology features, we have evaluated common SpA-related comorbid pathologies. By examining intestine, eyes and skin tissue during disease progression we did not observe any signs of pathology. However, we demonstrated with functional and immunohistochemical data a comorbid left-sided heart valve pathology confirming and expanding previous findings (21). More specifically, we observed that at 40 weeks of age, TgA86 mice displayed aortic valve thickening, consisting mainly of vimentin-positive mesenchymal cells and sparse Gr1-positive neutrophils (Additional file 1; Figure S1a, b). The aortic valve thickening and fibrosis resulted in functional changes including increased aortic velocity (Additional file 1; Figure S1c) with mild regurgitation which was observed in 10-20% of the tested 40-week old mice. The mitral valve was not affected (Additional file 1; Figure S1b), and hence mitral velocity remained unchanged (Additional file 1; Figure S1d). Importantly, ejection fraction was significantly decreased (Additional file 1; Figure S1e), indicating cardiac function impairment probably caused by aortic insufficiency. Despite their impaired heart valve function, TgA86 mice did not display premature death and their electrocardiogram analysis was normal.

### TgA86 axial and peripheral pathologies evolve through an early phase of inflammation and bone erosion gradually leading to new bone formation

The main pathological features as well as the progression of the axial pathology of TgA86 mice was further studied in the tail vertebrae that exhibit consistently the more pronounced pathology. As pathology first becomes apparent in the vertebrae proximal to the body of the mouse, all further analysis was based on evaluation of the first five-six caudal vertebrae.

Histopathological evaluation of H&E stained sections of tail vertebrae revealed signs of inflammation starting from 4 weeks of age and progressively worsening with time. By 20 weeks of age, 100% of the transgenic mice examined exhibited aggravated pathological features (Fig. 3a; H&E) at the intervertebral joint area with signs of severe enthesitis (Fig. 3a; H&E; black arrows) and intervertebral disk degeneration (Fig. 3a; H&E). These pathological features were homogeneous in the different vertebrae of the same mouse when examined at 20 weeks of age, but, as disease progressed, heterogeneity of inflammation between different animals and also between the different vertebrae of the same animal, was evident. Interestingly, while the percentage of vertebrae with high inflammation was increased at the age of 20 weeks in TgA86 mice, at the age of 40 weeks it declined with a concomitant two-fold increase in the percentage of vertebrae with low inflammation (Fig. 3c).

**Figure 3.**
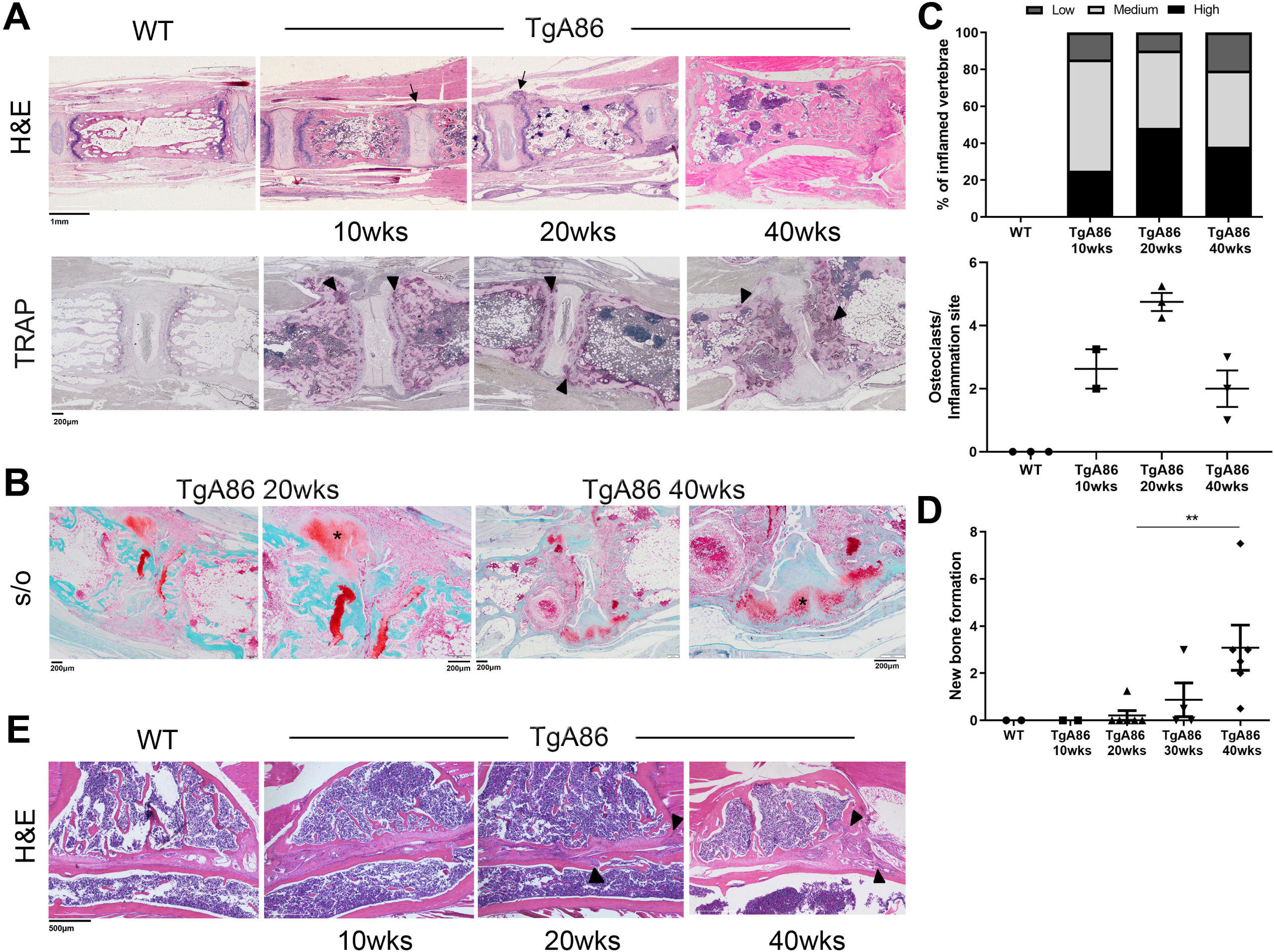
TgA86 axial pathology progresses through sequential stages of inflammation, bone erosion and ectopic new bone formation. **(A)** Representative H&E-stained sections of the tail vertebrae show enthesitis with inflammatory cell accumulation at the periphery of the intervertebral discs (black arrows) of TgA86 mice, while representative TRAP-stained tail vertebrae show the accumulation of active osteoclasts at the sites of inflammation (black arrow heads). **(B)** Representative safranin-O stained sections of tail vertebra show fibrocartilage formation at sites of enthesis in 20-week-old TgA86 mice and ectopic chondrocytes in 40-week-old TgA86 mice (black asterisk in higher magnification panels). **(C)** Inflammation is depicted as the percentage of vertebrae with low, intermediate or high scores. An increase in the percentage of joints with high inflammation is observed at 20-week-old TgA86 mice and this percentage decreases by 40 weeks of age. Similarly, there is an increase in the number of osteoclasts at 20 weeks of age that drops by 40 weeks of age. **(D)** Safranin-O staining of tail vertebra sections indicates increased presence of fibrocartilage (20 weeks) and of ectopic chondrocytes at later stages, 30 and 40 weeks of age. **(E)** Representative H&E-stained sections show inflammatory infiltrates accumulation in the sacroiliac joints of TgA86 mice (black arrowheads).

Additional pathological features that were observed as early as 10 weeks of age and progressively became more pronounced, included the roughening of the vertebral surface, the squaring of the vertebral body as well as alterations in the bone marrow cavity, where the formation of ectopic cartilaginous matrix and bone marrow cell aggregates was observed (Fig. 3a; H&E).

The extend of bone erosion was evaluated based on the number of osteoclasts stained with TRAP in tail sections of TgA86 mice at different ages. High numbers of osteoclasts were observed in the vertebral body of the tail of TgA86 mice highlighting an active bone remodeling process occurring in these mice (Fig. 3a; black arrowheads). More interestingly, when focusing on the osteoclasts present at the sites of inflammation (enthesitis), we observed an increase in their numbers between 10 and 20 weeks of age and a subsequent drop by 40 weeks of age (Fig. 3c).

Staining of TgA86 tail sections with Safranin-O allowed the detection of ectopic cartilage, gradually leading to new bone formation which is a main pathological feature of human SpA. Starting at 20 weeks of age, we could observe fibrocartilage at the inflamed edges of the vertebrae (enthesis) (Fig. 3b; black asterisk and Fig. 3d), while at later ages, 30 and 40 weeks of age, specifically at sites where inflammation was reduced or absent, we could detect the presence of ectopic chondrocytes that occasionally bridged adjacent vertebrae (Fig. 3b; black asterisk and Fig. 3d).

Overall, the axial pathology in TgA86 mice appeared to evolve through an initial phase that involved pronounced inflammation and bone erosion, while as the disease progressed, between 20 and 30 weeks of age, the signs of inflammation and bone erosion were reduced and replaced by the formation of new cartilaginous tissue, eventually leading to bridging of the vertebrae detected later on in disease. As sacroiliac joints are sites of preference for the manifestation of human SpA, we assessed histopathologically inflammation and tissue damage in the sacroiliac joints of TgA86 mice. As early as 10 weeks of age, TgA86 exhibited pannus-like tissue at the sacroiliac articular surface, while by 20 weeks of age, invasion of inflammatory cells to the sacrum and iliac bones was also evident (Fig. 3e; black arrowheads).

Peripheral pathology was studied by histopathological examination of the hind joints of TgA86 mice and was characterized by synovitis, enthesitis and focal subchondral bone erosions. By 10 weeks of age, TgA86 mice exhibited extensive enthesitis (Fig. 4a; arrow 1), pannus formation (Fig. 4a; arrow 2), and bone erosion (Fig. 4a; box), while by 20 weeks of age, pannus evolved to progressive destruction of the articular cartilage and subchondral bone (Fig. 4a; black asterisk) that, by 40 weeks of age, was greatly exacerbated.

**Figure 4.**
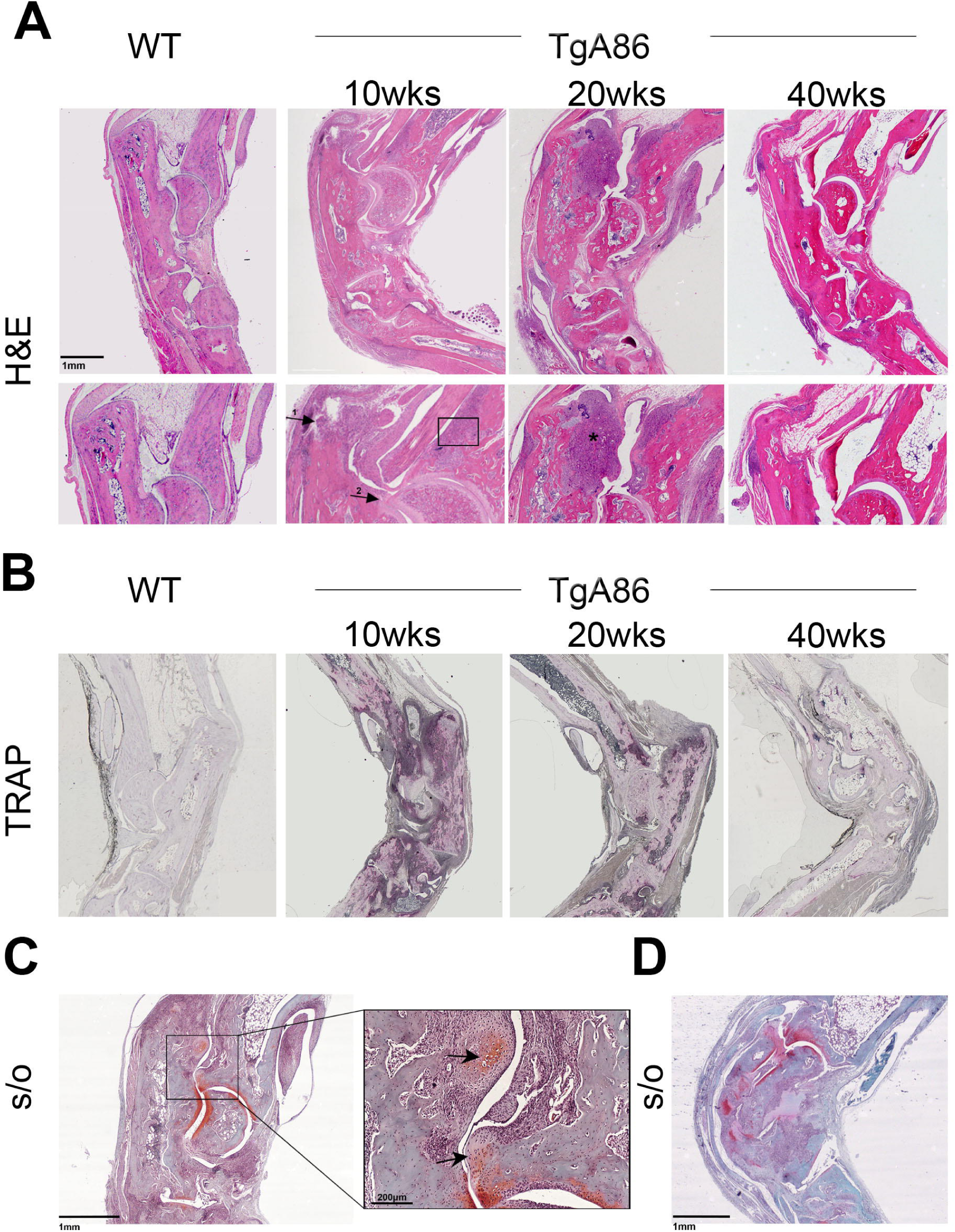
TgA86 peripheral disease progression involves features of inflammation, bone erosion and ectopic new bone formation. **(A)** Representative H&E stained sections of hind limbs show the accumulation of cells in the joints of TgA86 mice including features of enthesitis (arrow 1), synovitis (arrow 2), bone erosion (box) and bone destruction (asterisk) highlighted in higher magnification (lower panels). **(B)** Representative TRAP-stained sections of hind limbs show the accumulation of active osteoclasts in the joints of 10 and 20-week-old TgA86 mice, while their number is reduced in the joints of 40-week-old TgA86 mice. **(C, D)** Safranin O-stained sections of hind limbs of 20-week-old **(C)** and 40-week-old **(D)** TgA86 mice show the presence of fibrocartilage and ectopic chondrocytes (black arrows).

Increased numbers of osteoclasts were detected in the hind joints at 10 and 20 weeks of age by TRAP staining which however, similarly to our observations in the tail vertebrae, decreased at 40 weeks of age (Fig. 4b). Interestingly, µCT trabecular analysis of femurs of 20 and 40-week-old TgA86 mice revealed an osteoporotic phenotype further indicating progressive systemic bone loss (Additional file 1; Figure S2).

At 20 weeks of age, we also observed the formation of ectopic safranin-O stained cartilaginous tissue at the inflamed enthesis (Fig. 4c; arrows) that was more pronounced at 40 weeks of age (Fig. 4d), leading to the development of new ectopic bone tissue (Fig. 4c; arrows and Fig. 4d).

### New bone formation in tmTNF transgenic mice involves mechanisms of endochondral and membranous ossification

Immunohistochemical characterization of the axial pathology of 20-week-old TgA86 mice revealed that the increased cellularity observed during the inflammatory phase, consisted mainly of increased number of mesenchymal cells and neutrophils. More specifically, we observed accumulation of vimentin-positive mesenchymal cells at the enthesis and along the ligaments surrounding the vertebrae, but also inside the bone marrow cavity (Fig. 5a). Osteopontin staining at sites of cortical and cancellous bone as well as inside the bone marrow cavity appeared also increased, suggesting the accumulation of osteoblastic lineage cells, while increased periostin staining was also observed in affected vertebrae, and particularly at sites where bone erosion was evident, also suggesting increased osteoblastic activity (Fig. 5a).

**Figure 5.**
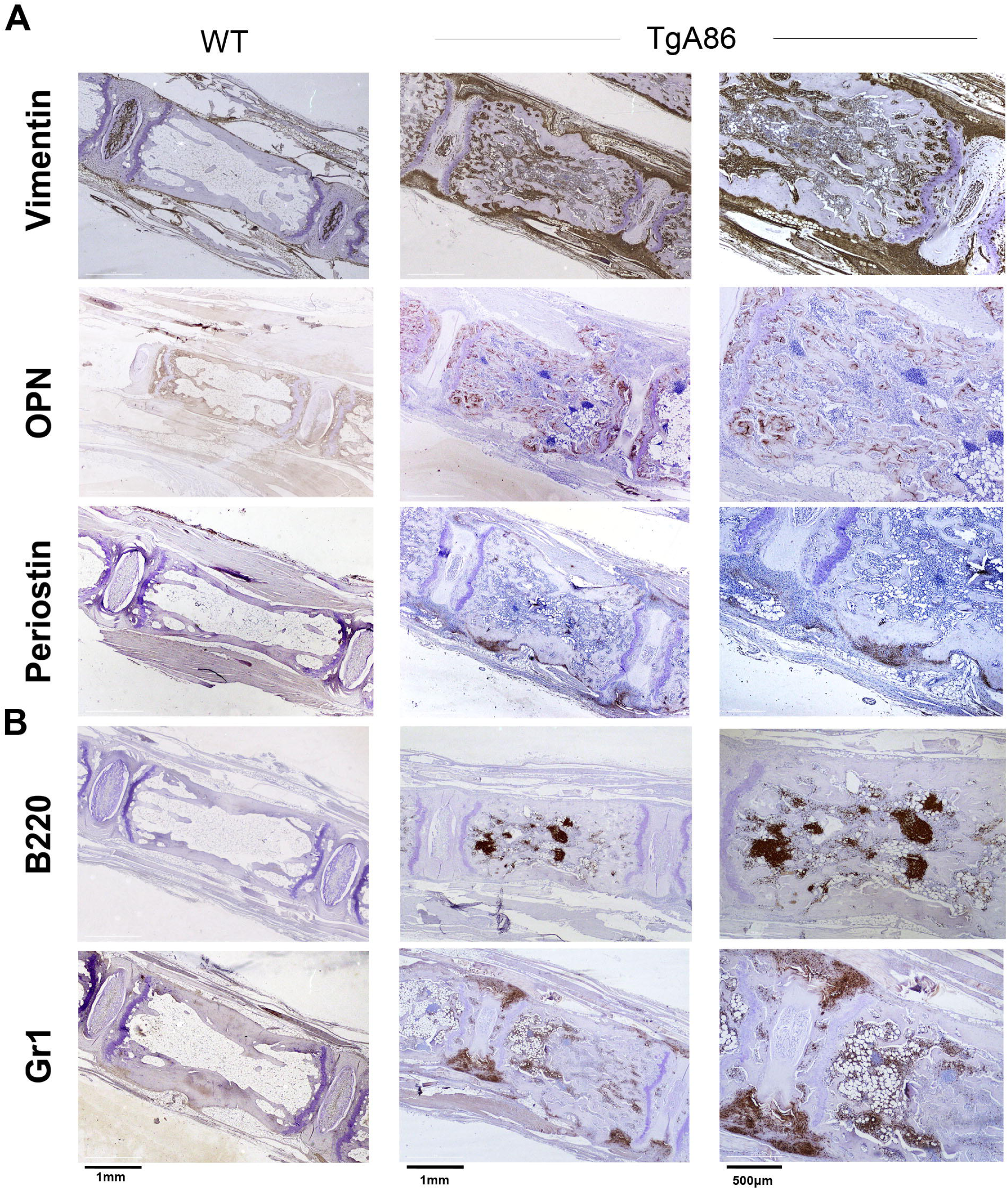
The cellular composition of the TgA86 axial pathology involves mesenchymal and osteoblastic lineage cells. **(A)** Vimentin, osteopontin (OPN) and periostin staining on the surface and the bone marrow cavity of the tail vertebral bodies as well as in the periphery of the intervertebral discs of 20-week-old TgA86 mice, indicate active new bone formation processes involving mesenchymal and osteoblastic lineage cells. **(B)** B220 positive cells form aggregates in the bone marrow cavity, while Gr1 positive cells accumulate at the sites of enthesis of 20-week-old TgA86 mice.

These stainings provide us with insights on the mechanisms that eventually lead to new bone formation observed at the advanced ages of TgA86 mice. More specifically, the increased vimentin and periostin staining indicates mesenchymal cell accumulation and increased osteoblastic activity in the periosteum that suggests an ongoing active membranous ossification mechanism. Similarly, the increased vimentin and osteopontin staining in the osseous vertebral cavity indicates vimentin-positive mesenchymal cell accumulation and condensation leading to osteopontin-positive osteoblastic lineage differentiation, suggesting an active endochondral ossification process.

Cell aggregates in the bone marrow cavity of TgA86 mice were found to consist mostly of B220-positive B cells, while Gr1-positive neutrophils accumulated mainly at the sites of the enthesis and were also loosely scattered in the bone marrow cavity and along the periosteum (Fig. 5b).

In a similar fashion to the axial pathology, peripheral pathology in the hind joints was also characterized by the accumulation of vimentin-positive mesenchymal cells and Gr1-positive neutrophils at the sites of synovitis and enthesitis, while there was only sparse presence of B220-positive B cells in the bone marrow cavity (Additional file 1; Figure S3).

### Anti-TNF therapy efficiently ameliorates both axial and peripheral TgA86 pathologies

We further assessed the response of the TgA86 model to anti-TNF treatment, a therapy that has proven efficacious in the early treatment of human SpA patients (22,23). More specifically, we treated TgA86 mice with Etanercept starting either from 2.5 weeks of age, i.e. early in the inflammation phase, or from 9 weeks of age, i.e. late in pathology when inflammation is more extensive leading to activation of cell types contributing to bone remodeling. Treatment continued up to 20 weeks of age and its effect was evaluated both *in vivo* and *ex vivo*.

Clinical monitoring of the TgA86 pathology was performed by assessing key phenotypic features of the disease evident up to 20 weeks of age. More specifically, peripheral pathology was assessed on a scale from 0 to 2 taking into consideration the severity of hind joint swelling, finger and limb deformation as well as grip strength (Additional file 1; Table S1), while axial pathology was assessed on a scale from 0 to 3 taking into consideration the tail pathology, including the number and extend of tail bendings as well as tail ankylosis (Additional file 1; Table S2). Early treatment with Etanercept completely abolished both the axial and peripheral clinical manifestations of the TgA86 pathology, while late treatment only partially ameliorated the peripheral arthritis pathology and had a minimal effect on the severity of the clinical manifestations of the axial pathology (Fig. 6a).

**Figure 6.**
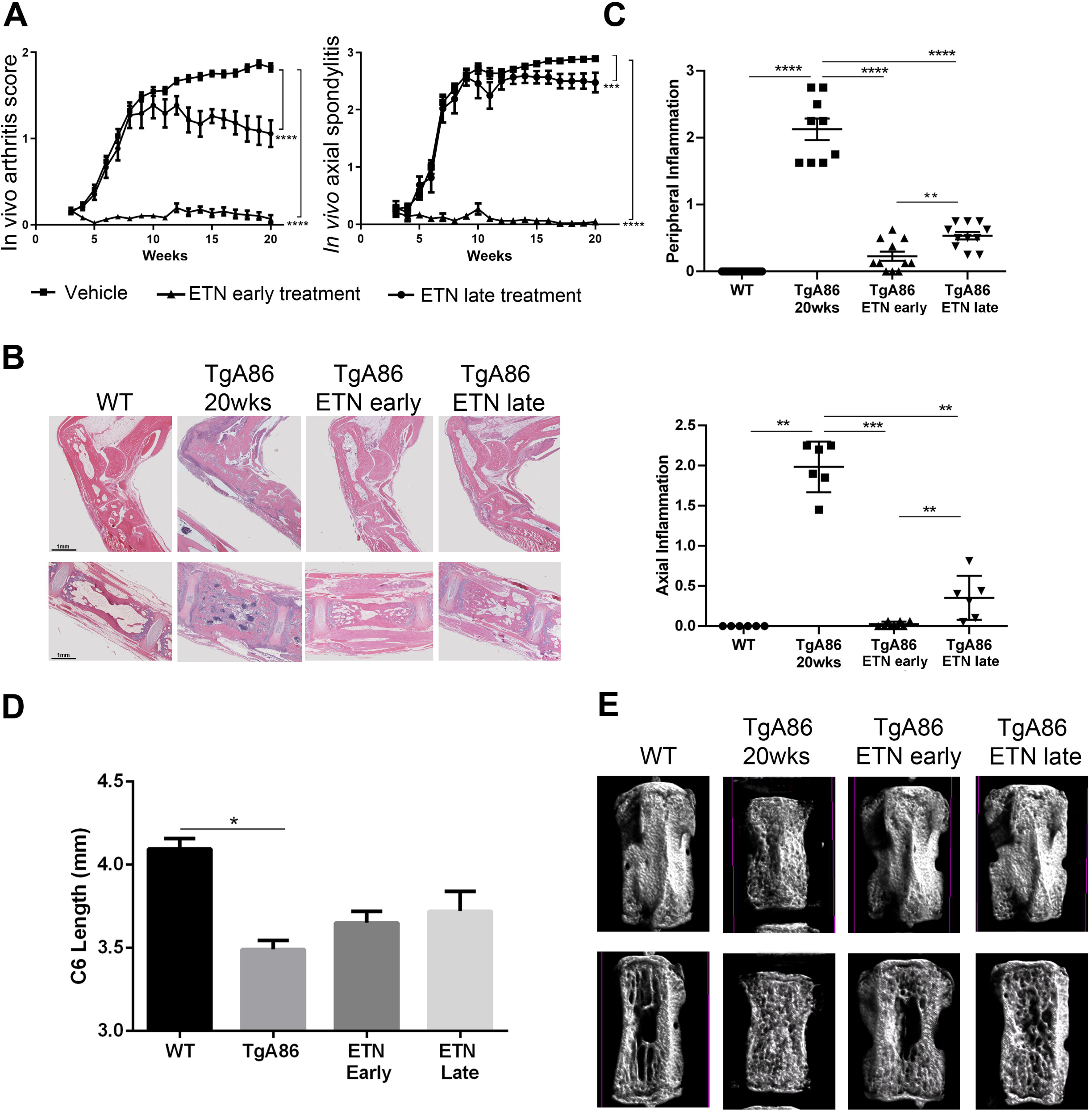
Early and late anti-TNF treatment can inhibit both the peripheral and axial TgA86 pathologies. **(A)** Early treatment with anti-TNF, starting at 2.5 weeks of age, and, to a lesser degree, late treatment, starting at 9 weeks of age, can ameliorate the clinical symptoms of both the peripheral and axial pathologies of 20-week-old TgA86 mice. **(B)** Representative H&E stained sections of hind limbs (top panel) and tail vertebrae (lower panel) show that both early and late anti-TNF treatment can ameliorate features of inflammation, bone erosion and bone marrow cell aggregates in 20-week-old TgA86 mice. **(C)** Both peripheral and axial inflammation of 20-week-old TgA86 mice is ameliorated following either early or late treatment with anti-TNF. Early anti-TNF treatment has statistically significant inhibitory effect on the TgA86 inflammation compared to late treatment. **(D)** Both early and late anti-TNF treatment can partially restore the reduced vertebral length as this is measured by μCT imaging of the 6^th^ caudal vertebra (C6) of 20-week-old TgA86 mice. **(E)** 3D μCT models (upper panel) and the longitudinal sections (lower panel) of reconstructed data sets of the C6 caudal vertebra of 20-week-old mice reveals that both early and late anti-TNF treatment can restore the surface, size and shape characteristics as well as the bone marrow cavity structure of TgA86 vertebrae to more closely resembling those of the wild type mice. [ETN: etanercept; Data are presented as mean±SEM; *P<0.0222; **P<0.004; ***P<0.0007; ****P<0.0001]

The effect of the early and late treatment protocols was also assessed histopathologically as well as by µCT analysis upon completion of the treatment. Histopathological evaluation of the axial and peripheral pathology involved the assessment of inflammation, as at 20 weeks of age this was the most prominent feature of the pathology. Axial inflammation was assessed in the first 5-6 caudal vertebra starting from the base of the tail, while peripheral inflammation was assessed in at least 3 different joints (ankle and metatarsal in sagittal plane) using a scoring scale from 0-3 (Additional file 1; Tables S3 and S4). Early treatment with Etanercept completely abolished both the axial and peripheral inflammation, while late treatment also greatly ameliorated both the axial and peripheral inflammation although to a slightly less extend compared to the early treatment protocol (Fig. 6b and 6c).

The structural changes of the TgA86 tail vertebrae and their improvement following Etanercept treatment were also visualized using 3D µCT modeling. The TgA86 tail vertebrae exhibited reduced length compared to the vertebrae of the wt mice and interestingly, early anti-TNF treatment, and to a lesser degree late treatment, could partially restore this feature (Fig. 6d). Moreover, 3D modeling highlighted the changes in the shape and surface of the TgA86 tail vertebrae that appeared squared with a rough surface and a dense matrix structure inside the bone marrow cavity, characteristics that were respectively restored following both early and late anti-TNF treatment to the bar belled shape and smoother surface with absence of dense matrix in the bone marrow cavity observed in wt mice vertebrae (Fig. 6e).

## Discussion

SpA progresses through an early inflammatory phase affecting the sacroiliac joints, the spine and the peripheral joints (24) and advances to a phase of reduced inflammation characterized by the formation of ectopic bone. This new bone formation leads to eventual joint ankylosis (ankylosing spondylitis) and functional disability (1,5,6).

As our understanding of how such as complex disease initiates and also progresses to its different pathological stages remains limited, there is a great need for the identification of prognostic and diagnostic markers, as well as for novel therapeutic approaches managing also the post-inflammatory phase of disease.

Therefore, the development and further exploitation of well-characterized animal models that highly reproduce the pathological features of SpA in a similar progressive manner as the one observed in human patients is crucial, especially if such models overexpress key molecules known to be involved in the development of the human pathology. TNF is a molecule proven to play a key pathogenic role in SpA (22), as its blockade is currently an effective therapy applied in the clinic for early treatment of human patients (22,25). Thus, a TNF-dependent model reproducing the full spectrum of the SpA pathological characteristics would be an invaluable addition in the field. To this end, we perform here a detailed analysis and characterization of the transmembrane TNF overexpressing TgA86 transgenic mouse model, that has been previously shown to develop arthritis (20), and we show that it reproduces main features of human SpA.

Similar to human patients, TgA86 mice developed SpA-like disease through an initial inflammatory phase manifested as enthesitis and subsequent erosive changes characterized by mesenchymal cell accumulation and neutrophilic infiltration, mainly at sites of attachment of the spinal ligament to bone. Later in the development of the disease, the inflammatory infiltrate is reduced and excessive tissue formation ensues, intervertebral disc degeneration and destruction and ectopic mature and hypertrophic chondrocytes that expanded between adjacent vertebrae in the caudal area of the axial skeleton. This ectopic new bone formation led to vertebral fusion, as revealed by CT imaging analysis, resembling imaging findings detected in human patients of ankylosing spondylitis (26–28). Interestingly, severe inflammation was never found simultaneously with severe excessive tissue formation within the same joint supporting sequential, rather than parallel progression of these disease features. This led to a significant variability of the stage of the pathology seen at different segments of the skeleton of the same mouse, similar to what happens in human patients, as in one vertebra junction new bone formation could be in progress, whereas the next one might still be at the inflammatory phase. Finally, sacroiliitis was also observed in the TgA86 mice involving mainly synovitis and enthesitis, however to a lesser degree than expected according to the human manifestations.

TgA86 mice developed all the features of human SpA, however with a distinct anatomical distribution to that of the human patients. It has been previously reported that skeletal mechanical stress and tissue microdamage are among the early events preceding the development of entheseal inflammation and the establishment of SpA in humans (24) but also in mice, where it was shown that hind limb unloading in the TNF *ΔARE* mice significantly suppressed inflammation of the Achilles tendon (29). Clearly, the biomechanical stress applied during ligament and tendon action differ significantly between species. Therefore, the obvious anatomical differences and distribution of pressure on the axial skeleton between mice and humans could explain the localization of the TgA86 axial pathology primarily at the sacral (S3-S4 junction) and caudal vertebrae that demonstrate increased movement in mice compared to humans, and the rather mild pathology observed in the sacroiliac joints which are the first sites affected in human patients (1).

As suggested by Benjamin *et al.* 2009 (30), entheseal new bone formation in humans follows a distinctive pattern involving all three ossification mechanisms, i.e. membranous, endochondral and chondroidal ossification, as is also the case in TgA86 mice. More specifically, loss of normal vertebral shape of TgA86 tails was associated with histopathological findings of mesenchymal cell expansion (vimentin positive cells) and osteoblast activation (periostin staining), both indicative of active intramembranous ossification. Additionally, the presence of a chondroid-osseous matrix in the bone marrow cavity, in combination with the accumulation of osteoblastic lineage cells (osteopontin staining), indicate an active endochondral ossification mechanism. Finally, mesenchymal expansion at the intervertebral disc areas and accumulation of mature and hypertrophic chondrocytes suggest an active chondroidal ossification mechanism resembling endochondral ossification mainly connected to ectopic bone formation (31).

Our findings support that the TgA86 model nicely reproduces cardinal features of the axial and peripheral pathology of the human SpA and, notably, we also show that these mice develop progressive systemic bone loss (osteoporosis), indicated by μCT trabecular analysis of the femur, as well as heart valvular dysfunction with both pathologies suggested to be prevalent comorbidities in human SpA patients (7,32,33). Overall, these mice could be an advantageous SpA mouse model as they spontaneously mimic the complexity of human SpA in a tm-TNF-driven manner. The main findings of the axial pathology, as well as the new bone formation observed in the TgA86 mice could differentiate this model from the other mouse or human TNF overexpressing mice (16–19,21) that express soluble and transmembrane TNF signaling through both TNFR1 and TNFR2 and have been reported to develop only peripheral arthritis. This could suggest that tmTNF-TNFR2 signaling may have a central role in the development of SpA, a hypothesis also supported by the high levels of tmTNF detected in human SpA patients (34).

Blockade of TNF early in the development of SpA has proven highly advantageous in human patients alleviating early symptoms and signs of the disease and, more importantly, precluding further development of structural lesions such as ankylosis (25). Similarly, we show here that both the peripheral as well as the axial pathology of the TgA86 model is ameliorated with anti-TNF treatment when this is initiated at an early stage of the disease during the inflammatory phase, when signs of new bone formation are absent. However, major challenges in the treatment of SpA remain, especially concerning patients at later stages of the disease, as early diagnosis is not always possible. The TgA86 model faithfully reproducing the two distinct phases of the SpA pathology allows the design of testing protocols both for the evaluation of drugs targeting the inflammatory as well as the post-inflammatory phase of the SpA pathology.

## Conclusion

Our work shows that the TgA86 tmTNF transgenic mouse model faithfully reproduces important features of human SpA, including the development of axial and peripheral pathology that progresses through an early inflammatory phase, with mesenchymal cell involvement, accompanied by bone erosion and advances to activation of new bone formation mechanisms and subsequent ankylosis, along with the manifestation of extraarticular heart valve pathology and osteoporosis. We believe that coexistence of all these aspects of the SpA pathology in one animal model render it an invaluable tool towards the study of the relevant importance of different pathways in disease initiation and progression, mechanisms of osteoproliferation, the role of entheseal resident cells as well as the role of TNF and other pivotal pathogenic cytokines in the development and progression of SpA. Moreover, the TgA86 SpA mouse model can serve as basis for the development of experimental protocols for the evaluation of the efficacy of therapeutics targeting both the early inflammatory and late osteoproliferative stages of SpA.

## List of abbreviations

AS: Ankylosing Spondylitis
CT: Computed Tomography
CVD: Cardiovascular Disease
EDTA: Ethylenediaminetetraacetic acid
H&E: Hematoxylin&Eosin
IBD: Inflammatory Bowel Disease
MRI: Magnetic Resonance Imaging
OP: Osteoporosis
OPN: Osteopontin
PsA: Psoriatic arthritis
RA: Rheumatoid Arthritis
ReA: Reactive arthritis
s/o: Safranin/O
SpA: Spondyloarthritis
Tg: transgenic
tmTNF: transmembrane mouse
TNF TNF-α: Tumor Necrosis Factor α
TRAP: Tartrate-resistant acid phosphatase
UTR: Untranslated Region
wt: Wild Type
µCT: micro CT

## Declarations

### Ethics approval and consent to participate

This study was carried out in accordance with the recommendations of Institutional Committee of Protocol Evaluation in conjunction with the Veterinary Service Management of the Hellenic Republic Prefecture of Attika (Approval No.141688 24/04/19) and in accordance to national legislation and the European Union Directive 63/2010.

### Consent for publication

Not applicable

### Availability of data and materials

All data that support the findings of this study are included in the article or uploaded as additional file.

### Competing interests

GK participates in the BoD of Biomedcode; FM has received grants from Pfizer. All other authors declare no competing interests.

### Funding

This work was co-financed by the European Union and Greek national funds through the Operational Program Competitiveness, Entrepreneurship and Innovation, under the call RESEARCH-CREATE-INNOVATE (Project code: T2EΔK-03807; SEPIA).

### Author’s Contributions

GK, ECV, NK and MCD designed the study and interpreted the experimented results. ECV, CG, LN, KK, EA, IM and MR performed the experiments and data analysis. FM, MA, CP, and GL contributed to data interpretation. EC, NK and MCD wrote the first draft of the manuscript and all authors were involved in critically revising its final manuscript. All authors read and approved the final manuscript.

## Acknowledgments

The authors thank Maria Mathopoulou for her help on processing and staining histological samples and Olga Graphou for lab and animal facility management.

Parts of this work have been presented at the 2017 EWRR Congress (Abstract 06.13; Athens, Greece, 2017) and at the 2018 EULAR Congress (Abstract FRI0163; Amsterdam, Netherlands, 2018).

## Additional file 1

**Table S1: Clinical evaluation of peripheral arthritis (hind limbs)**

**Table S2: Clinical evaluation of axial spondylitis (tail)**

**Table S3: Histopathological evaluation of peripheral inflammation (hind limbs)**

**Table S4: Histopathological evaluation of axial inflammation (per vertebra)**

**Table S5: Histopathological evaluation of new bone formation**

**Figure S1: Histopathological and functional analysis of the TgA86 hearts reveal comorbid aortic heart valve dysfunction. (A, B)** Representative immunohistochemical stainings of aortic valve sections of wt and TgA86 mice at the age of 20 weeks with anti-Vimentin and anti-Gr1 antibodies **(A)** reveal that the thickened areas of the valve **(B)** are mainly composed by Vimentin+ resident fibroblasts as well as few Gr1+ infiltrated neutrophils. **(C, D)** Blood aortic (AoV) and mitral (MV E and A) velocity measurements, acquired by Doppler analysis of TgA86 mice and wt littermates at 40 weeks of age, reveal increased AoV, indicating aortic valve stenosis, while the function of mitral valve remains unaffected (data are presented as individual values, with mean±SEM). (**E**) Reduced Ejection fraction (EF%) of TgA86 mice at 40 weeks of age, indicate contractile dysfunction of their left ventricle. (Data are presented as individual values, with mean±SEM; **P<0.01).

**Figure S2. TgA86 mice show enhanced bone resorption leading to osteoporosis-like pathology.** Trabecular analysis of the metapheseal regions of the femurs of 20 and 40-week-old TgA86 and wt mice by μCT show progressively reduced BV/TV (bone volume/ total volume, %), Tb.N (trabecular number per mm^-1^) and Con.Dens. (connectivity density, mm^-3^) as well as increased Tb.Sp (trabecular separation, mm).

**Figure S3. Immunohistochemical analysis of TgA86 peripheral pathology reveals Vimentin- and Gr1-positive cell accumulation at the sites of enthesis.** Vimentin and Gr1 positive cells accumulate at sites of enthesis at the hind joints of 20-week-old TgA86.

## Notes

### Competing Interest Statement

Prof. George Kollias participates in the BoD of Biomedcode; Dr. Florian Meier has received grants from Pfizer. All other authors declare no competing interests.

