## Supplementary file for "Ectopic bone formation and systemic bone loss in a transmembrane TNF-driven model of human spondyloarthritis"

### Additional file 1

- **Supplementary Tables**
- **Supplementary Figures**

**Table S1**

| CLINICAL EVALUATION OF PERIPHERAL ARTHRITIS (HIND LIMBS) | |
|---|---|
| **SCORE** | **CHARACTERISTICS** |
| 0 | No signs of arthritis |
| 1 | Mild to moderate arthritis (joint distortion by swelling, inflamed paw) |
| 2 | Moderate to severe arthritis (severe joint, paw and finger swelling, joint – leg deformation, no grip strength) |

**Table S2**

| CLINICAL EVALUATION OF AXIAL SPONDYLITIS (TAIL) | |
|---|---|
| **SCORE** | **CHARACTERISTICS** |
| 0 | Normal tail phenotype |
| 1 | Mild; Normal tail with first signs of tail bending in one or in several locations |
| 2 | Moderate to Severe; Multiple tail bendings |
| 3 | Severe; Tail stiffness with several tight tail bendings |

**Table S3**

| HISTOPATHOLOGICAL EVALUATION OF PERIPHERAL INFLAMMATION (HIND LIMBS) | |
|---|---|
| **SCORE** | **CHARACTERISTICS** |
| 0 / no disease | Normal Synovium; no inflammatory infiltrates |
| 1 / mild disease | Mild thickening of the synovial membrane; mild infiltration of inflammatory cells into periarticular tissue; inflammatory sites can be found on some but not all tarsal/ankle joints |
| 2 / moderate to severe | Increased thickening of the synovial membrane; enhanced infiltration of inflammatory cells; inflammatory sites are present on most but not all tarsal/ankle joints |
| 3 / severe disease | Massive accumulation of inflammatory cells throughout the whole specimen (synovial membrane, periarticular and connective tissue); severe hyperplasia of the synovial membrane, inflammatory tissue affects all tarsal/ankle joints |

**Table S4**

| HISTOPATHOLOGICAL EVALUATION OF AXIAL INFLAMMATION[1] (per vertebra) | |
|---|---|
| **SCORE** | **CHARACTERISTICS** |
| 0 / no disease | Healthy tissue, no inflammatory infiltrates |
| 1 / mild disease | Mild enthesitis; adjacent tissue not affected |
| 2 / moderate to severe | Moderate enthesitis and “pannus like” formation; signs of inflammation in adjacent tissues (tendons, muscle) |
| 3 / severe disease | Severe inflammation and “pannus like” formation that extends in tendon, muscle and intravertebral disc area |

[1] Final score of each mouse is calculated as the average of the first 4-6 caudal vertebral scores

**Table S5**

| HISTOPATHOLOGICAL EVALUATION OF NEW BONE FORMATION | |
|---|---|
| **SCORE** | **CHARACTERISTICS** |
| 0 | Normal |
| 1 | Signs of fibrocartilage formation in enthesis sites of each vertebra (pale or no s/o staining) |
| 2 | Ectopic chondrocytes in a small area around the intervertebral disc (s/o stain) |
| 3 | Ectopic chondrocytes in multiple areas around the intervertebral disc leading to fusion between vertebrae; hypertrophic chondrocytes are present (s/o stain) |

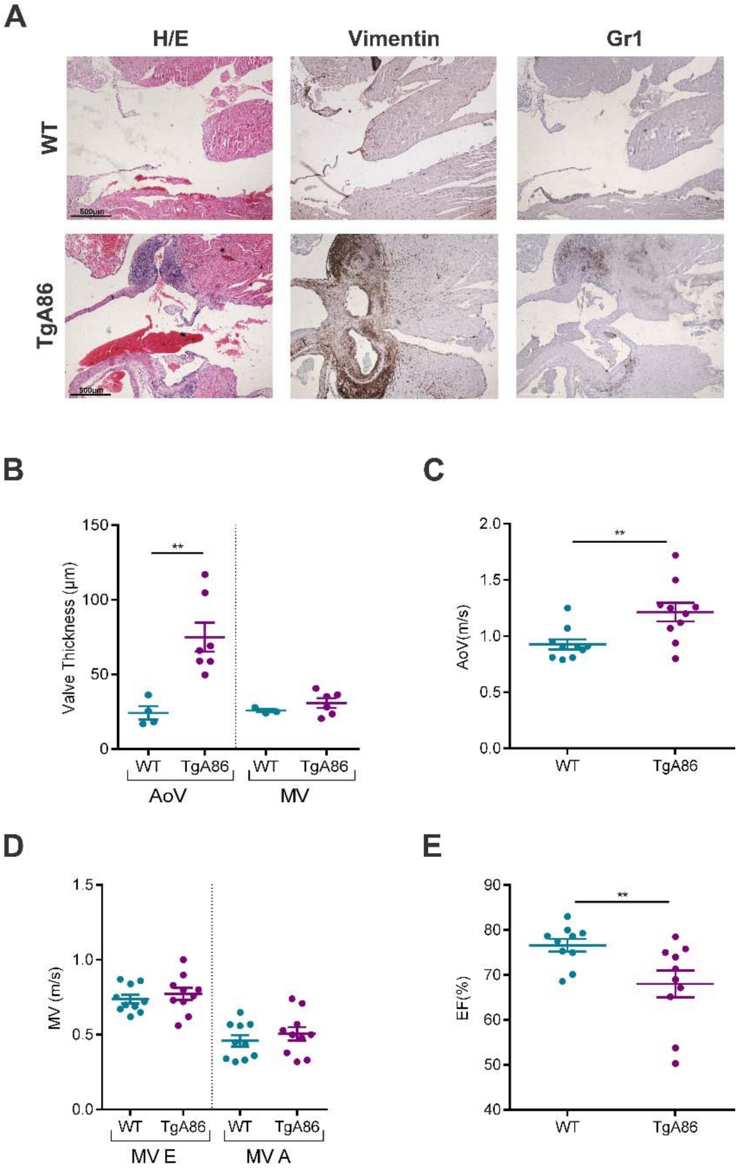


**Figure S1: Histopathological and functional analysis of the TgA86 hearts reveal comorbid aortic heart valve dysfunction. (A, B)** Representative immunohistochemical stainings of aortic valve sections of wt and TgA86 mice at the age of 20 weeks with anti-Vimentin and anti-Gr1 antibodies **(A)** reveal that the thickened areas of the valve **(B)** are mainly composed by Vimentin+ resident fibroblasts as well as few Gr1+ infiltrated neutrophils. **(C, D)** Blood aortic (AoV) and mitral (MV E and A) velocity measurements, acquired by Doppler analysis of TgA86 mice and wt littermates at 40 weeks of age, reveal increased AoV, indicating aortic valve stenosis, while the function of mitral valve remains unaffected (data are presented as individual values, with mean±SEM). (**E**) Reduced Ejection fraction (EF%) of TgA86 mice at 40 weeks of age, indicate contractile dysfunction of their left ventricle. (Data are presented as individual values, with mean±SEM; **P<0.01).

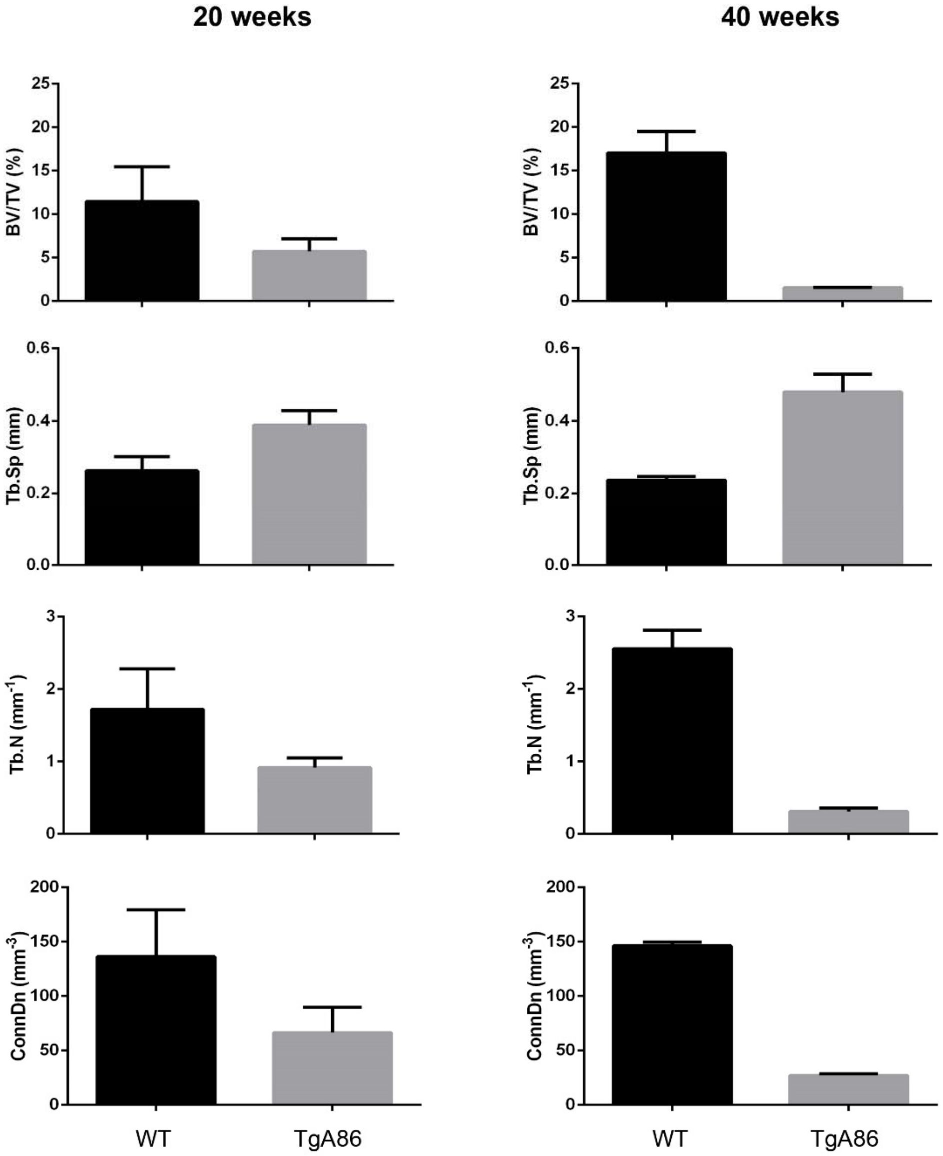


**Figure S2. TgA86 mice show enhanced bone resorption leading to osteoporosis-like pathology.** Trabecular analysis of the metapheseal regions of the femurs of 20 and 40 week old TgA86 and wt mice by µCT show progressively reduced BV/TV (bone volume/ total volume, %), Tb.N (trabecular number per $mm^{-1}$) and Con.Dens. (connectivity density, $mm^{-3}$) as well as increased Tb.Sp (trabecular separation, mm).

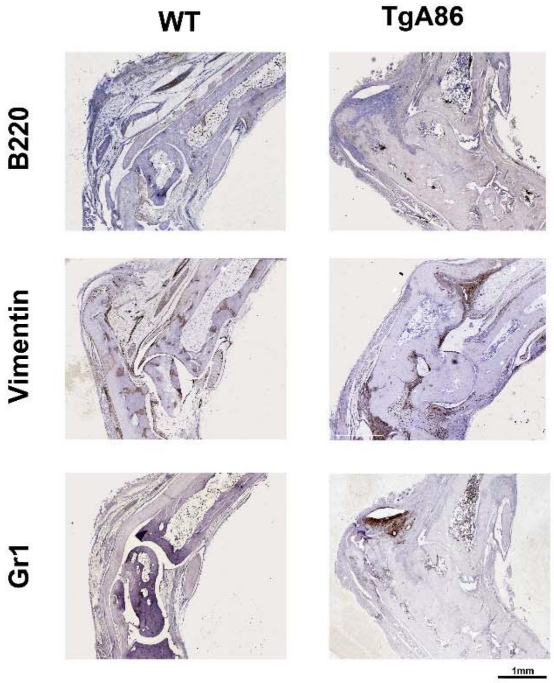


**Figure S3. Immunohistochemical analysis of TgA86 peripheral pathology reveals Vimentin- and Gr1-positive cell accumulation at the sites of enthesis.**

Vimentin and Gr1 positive cells accumulate at sites of enthesis at the hind joints of 20-week-old TgA86.
